# Helminth infection-induced carcinogenesis: spectrometric insights from the liver flukes, *Opisthorchis* and *Fasciola*

**DOI:** 10.1101/606772

**Authors:** Maria João Gouveia, Maria Y. Pakharukova, Banchob Sripa, Gabriel Rinaldi, Paul J. Brindley, Viatcheslav A. Mordvinov, Fátima Gärtner, José M. C. da Costa, Nuno Vale

## Abstract

Chronic infections with the flatworm parasites *Opisthorchis viverrini, Clonorchis sinensis* and *Schistosoma haematobium* are classified as group 1 biological carcinogens, i.e. definitive causes of cancer. In addition, we reported findings that support the inclusion of *Opisthorchis felineus* in this list of biological carcinogens. By contrast, infections with close phylogenetic relatives including *Fasciola hepatica* have not been associated with carcinogenesis. Earlier reports revealed of oxysterol metabolites of *Opisthorchis* liver fluke origin conjugated with DNA bases, suggesting that the generation of these DNA-adducts may underlie the mutagenicity and carcinogenicity of the infection with these food-borne pathogens. Here we employed liquid chromatography-mass spectrometry (LC-MS/MS) to investigate, compare and contrast spectrograms of soluble extracts from *F. hepatica* adult worms from bile ducts of cattle with those from *O. viverrini* and *O. felineus* from experimentally-infected hamsters. *F. hepatica* displayed a complex spectrophotometric profile. *F. hepatica* and *Opisthorchis* spp. shared several common compounds including oxysterol-like metabolites, bile acids and DNA-adducts, but the spectrometric profiles of these *Opisthorchis* species included far fewer compounds than *F. hepatica*. These findings support the postulate that oxysterol-like metabolites of parasite origin can initiate carcinogenesis and they point to a molecular basis for the inconsistencies among major groups of liver flukes concerning infection-induced malignancy.

**Author Summary:** Several species of trematodes are parasites of the human hepatobiliary tract. Infection with two of these flukes, *Clonorchis sinsensis* and *Opisthorchis viverrini*, fresh water fish-borne parasites that occur in East Asia is classified as group 1 carcinogens by the International Agency for Research on Cancer (IARC), i.e. definitive causes of cancer in humans. By contrast, infection with a different liver fluke, *Fasciola hepatica*, does not lead to malignant transformation of the biliary tract. Given the close phylogeny of all three parasites, this difference in carcinogenicity is intriguing and, if explained, likely of value in novel therapeutic approaches. The importance of the current findings is informative because they present a mass spectrometric analysis and catalog of the similarities and differences between fluke of the genus *Opisthorchis* and *F. hepatica*, potentially identifying carcinogenic metabolites of liver fluke origin. These metabolites can be expected to provide deeper understanding of helminth infection induced malignancy.

## Introduction

More than 20% of cancer in the developing world are caused by infections [1]. The World Health Organization’s International Agency for Research on Cancer (IARC) recognizes the infection with about 12 pathogens as group 1 biological carcinogens, i.e., definitive causes of cancer. These group 1 agents include three helminth parasites, specifically the fish-borne trematodes (FZT) *Opisthorchis viverrini* and *Clonorchis sinensis* and the blood fluke, *Schistosoma haematobium* [2]. In addition, we reported findings from hamster infection that support the inclusion of *Opisthorchis felineus*, also an FZT, to this list of biological carcinogens and definitive cause of cholangiocarcinoma [3]. We hypothesised that these helminths produce and release derivatives of oestrogens and oxysterols that promote oxidation of host DNA, inducing lesions, adducts and mutations [1,3–6]. The findings supported the postulate that these infection-associated cancers originate from a biological and/ or chemical insult followed by chronic inflammation, fibrosis, and a change in the tissue microenvironment that leads to a pre-cancerous niche [7,8]. Paradoxically, infections with other close phylogenetic relatives of these carcinogenic helminths, also food borne trematodes of the *Phylum Platyhelminthes* (Table 1), have not been categorized as group 1 biological carcinogens [9–15].

**Table 1.**
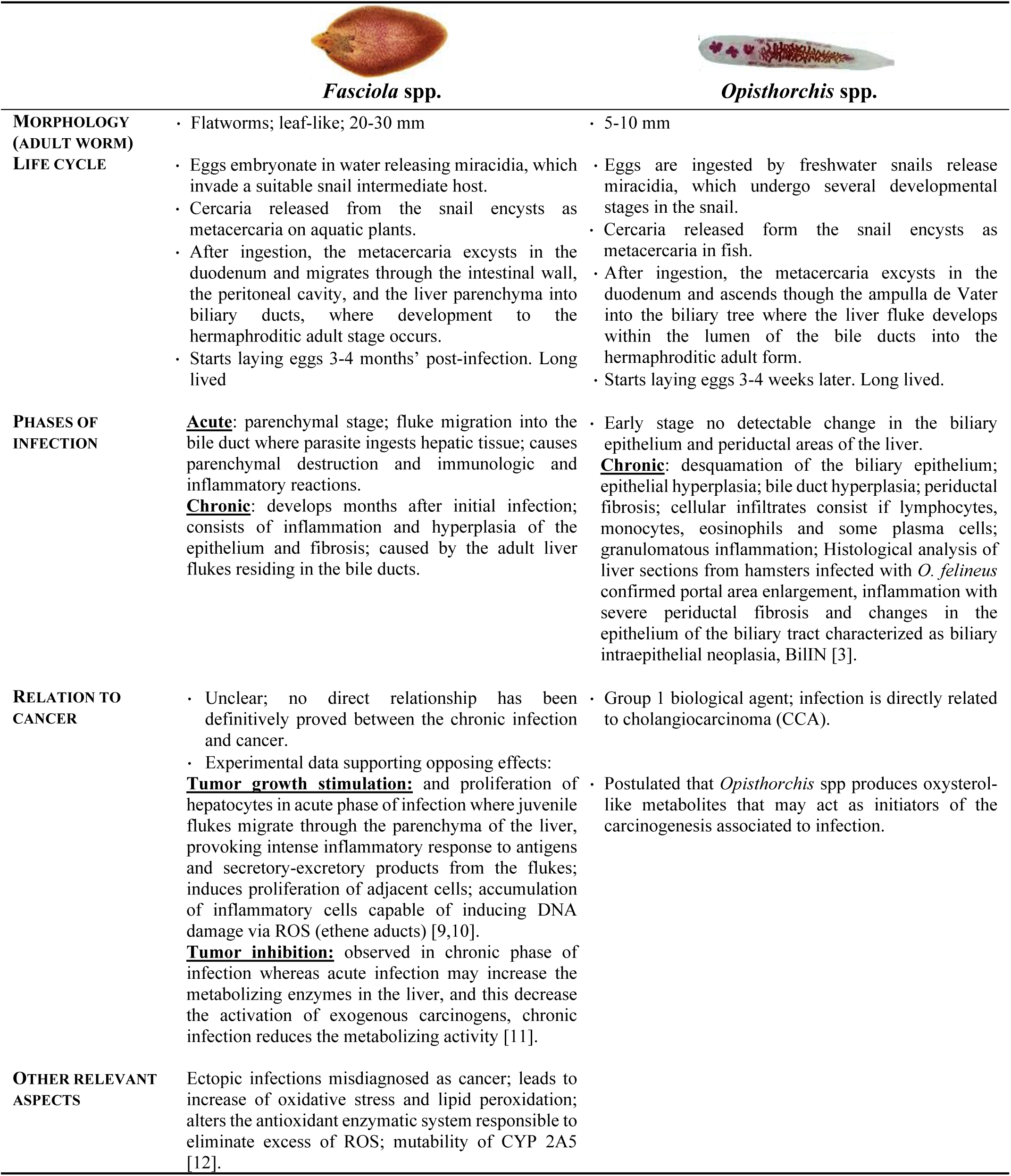
Comparison of morphology, life cycle and pathogenesis between *Fasciola hepatica* and *Opisthorchis* species.

|  | *Fasciola spp.* | *Opisthorchis spp.* |
| --- | --- | --- |
| **MORPHOLOGY (ADULT WORM)** | • Flatworms; leaf-like; 20-30 mm | • 5-10 mm |
| **LIFE CYCLE** | • Eggs embryonate in water releasing miracidia, which invade a suitable snail intermediate host.<br>• Cercaria released from the snail encysts as metacercaria on aquatic plants.<br>• After ingestion, the metacercaria excysts in the duodenum and migrates through the intestinal wall, the peritoneal cavity, and the liver parenchyma into biliary ducts, where development to the hermaphroditic adult stage occurs.<br>• Starts laying eggs 3-4 months' post-infection. Long lived | • Eggs are ingested by freshwater snails release miracidia, which undergo several developmental stages in the snail.<br>• Cercaria released from the snail encysts as metacercaria in fish.<br>• After ingestion, the metacercaria excysts in the duodenum and ascends through the ampulla de Vater into the biliary tree where the liver fluke develops within the lumen of the bile ducts into the hermaphroditic adult form.<br>• Starts laying eggs 3-4 weeks later. Long lived. |
| **PHASES OF INFECTION** | **Acute:** parenchymal stage; fluke migration into the bile duct where parasite ingests hepatic tissue; causes parenchymal destruction and immunologic and inflammatory reactions.<br>**Chronic:** develops months after initial infection; consists of inflammation and hyperplasia of the epithelium and fibrosis; caused by the adult liver flukes residing in the bile ducts. | • Early stage no detectable change in the biliary epithelium and periductal areas of the liver.<br>• **Chronic:** desquamation of the biliary epithelium; epithelial hyperplasia; bile duct hyperplasia; periductal fibrosis; cellular infiltrates consist of lymphocytes, monocytes, eosinophils and some plasma cells; granulomatous inflammation; Histological analysis of liver sections from hamsters infected with *O. felinus* confirmed portal area enlargement, inflammation with severe periductal fibrosis and changes in the epithelium of the biliary tract characterized as biliary intraepithelial neoplasia, BilIN [3]. |
| **RELATION TO CANCER** | • Unclear; no direct relationship has been definitively proved between the chronic infection and cancer.<br>• Experimental data supporting opposing effects:<br>**Tumor growth stimulation:** and proliferation of hepatocytes in acute phase of infection where juvenile flukes migrate through the parenchyma of the liver, provoking intense inflammatory response to antigens and secretory-excretory products from the flukes; induces proliferation of adjacent cells; accumulation of inflammatory cells capable of inducing DNA damage via ROS (ethene aducts) [9,10].<br>**Tumor inhibition:** observed in chronic phase of infection whereas acute infection may increase the metabolizing enzymes in the liver, and this decrease the activation of exogenous carcinogens, chronic infection reduces the metabolizing activity [11]. | • Group 1 biological agent; infection is directly related to cholangiocarcinoma (CCA).<br>• Postulated that *Opisthorchis* spp produces oxysterol-like metabolites that may act as initiators of the carcinogenesis associated to infection. |
| **OTHER RELEVANT ASPECTS** | Ectopic infections misdiagnosed as cancer; leads to increase of oxidative stress and lipid peroxidation; alters the antioxidant enzymatic system responsible to eliminate excess of ROS; mutability of CYP 2A5 [12]. |  |

For instance, *Fasciola hepatica* has a wide geographical range, causes major economic loss in sheep and cattle worldwide, and also is an important food borne trematodes (FBT) pathogen of humans [16]. Fascioliasis can induce host DNA damage through action of reactive nitric species (RNS) or oxygen species (ROS) [15,17]. We hypothesized that the first step for carcinogenesis associated with the chronic infection with *S. haematobium* and *O. viverrini* involved the ability of parasite metabolites to directly promote DNA lesions [5,6]. Seeking new insights in the apparent paradox of differences in carcinogenicity among closely related FBZ, we have analyzed extracts of adult worms of *F. hepatica, O. viverrini and O. felineus* using liquid chromatography coupled with mass spectrometry (LC-MS/MS). Remarkably, the LC-MS/MS chromatograms for each liver fluke species exhibited clear differences in regard the presence of oxysterols. These metabolites were minor components of the extract from *F. hepatica*, in contrast to the abundance and diversity of forms of oxysterols in *O. viverrini and O. felineus*. The presence of abundant oxysterols in the metabolites of *Opisthorchis* liver flukes support the notion that represent initiators of liver fluke infection-induced biliary tract malignancy.

## Material and methods

### Ethics Statement

Procedures undertaken complied with The Code of Ethics of the World Medical Association (Declaration of Helsinki) for animal experiments http://ec.europa.eu/environment/chemicals/lab_animals/legislation_en.htm. Syrian hamsters (*Mesocricetus auratus*) were purchased from the stock of the Puschino Animal Facility (Russia) and bred at the Animal Facility of the ICG SB RAS (RFMEFI61914X0005) (Russia). The hamsters were maintained according to protocols approved by the Committee on the Ethics of Animal Experiments of the Institute of Cytology and Genetics (Permit Number: 25 of 12.12.2014).

### Soluble extracts from *F. hepatica, O. viverrini* and *O. felineus* adult liver flukes

Adult worms of *F. hepatica* were obtained from the bile ducts of infected cattle at a local slaughterhouse [18]. It should be noted that the animals were processed as part of normal work of the slaughterhouse. *O. viverrini* and *O. felineus* were obtained as previously described [3,4]. In brief, metacercariae of *Opisthorchis* species were obtained from naturally infected cyprinoid fish in Khon Kaen province, Thailand or from naturally infected fish (*Leuciscus idus*) in the Ob River near the city of Novosibirsk, Siberia Russia, respectively. The fish were digested with pepsin-HCl [3]. Fifty metacercariae were used to infect hamsters (*Mesocricetus auratus*) and three months after infection, the animals were euthanized and adult *O. viverrini* or *O. felineus* flukes recovered from their bile ducts. The worms were washed extensively in phosphate buffered saline (PBS, pH 7.4) supplemented with 100 μg/mL streptomycin and 100 U/mL penicillin G and cultured overnight in serum free RPMI-1640 medium (Lonza, Basel, Switzerland) containing 1% glucose, and protease inhibitors (0.1 mM phenylmethanesulfonyl fluoride, 2 μM E-64 and 10μM leupeptin) (Sigma-Aldrich, St. Louis, Missouri) at 37 °C, 5% CO_2_.

Soluble extracts from all samples were prepared by sonication (5 × 5s burst, output cycle 4, Branson Sonifier 450, Germany) in PBS supplemented with protease inhibitors [500 μM 4-(2- aminoethyl) benzenesulfonyl fluoride hydrochloride (AEBSF), 0.3 μM aprotinin, 10 μM E-64, 10 μM bestatin and 10 μM leupeptin] (M221, Amresco, Solon, OH, USA), followed by 30 min centrifugation at 10,000 rpm, 4 °C. The protein concentration of supernatants was determined using a commercial kit. Ascorbic acid was added to 1 mg/ml to these extracts, which were stored in aliquots at −80 °C [3,4].

### Sample preparation and LC-MS/MS analysis

Samples were prepared and processed using liquid chromatography diode array detection electron spray ionization mass spectrometry, as described [3–5]. Due to the acceptable chromatographic performance of methanol as the solvent in terms of separation and sensitivity, with short gradient times [19], this solvent was added up to 20% (v/v). High performance liquid chromatography coupled with mass spectrometer was employed to investigate molecular species from liver flukes, with samples of 25 μL injected into the LC-MS/MS instrument for analysis. The mass analysis was performed within an LTQ Orbitrap XL mass spectrometer (Thermo Fischer Scientific, Bremen, Germany), fitted with an ultraviolet (UV) photo diode array (PDA) detector. Analysis involved a Macherey-Nagel Nucleosil C18- column (250 mm × 4 mm internal diameter; 5 μm particle diameter, end-capped), proceeding at a flow rate of 0.3 ml/min. The capillary voltage of the electrospray ionization was 28 kW, capillary temperature was 310 °C, flow rates of the sheath gas and auxiliary N2 were set to 40 and 10 (arbitrary unit as provided by the software settings), respectively, and gas temperature was 275 °C [3–5]. The mobile phase consisted of 1% formic acid in water (A)/acetonitrile (B) mixtures. Eluates were monitored for 75 min, run with a mobile phase gradient of 0-5 min, 100% A; 5-10 min, linear gradient from 100% to 80% A, 10-15 min 80% A, 15-50 min, linear gradient from 80% to 40% A; 50-65 min, 40% A; 65-75 min, linear gradient from 40% to 100% B. Washing for 15 min with acetonitrile was carried out to stabilize the column. Data were collected in negative electrospray ionization negative mode scanning a mass to charge ratio (m/z) range of 50-2,000.

## Results

### Both species of Opisthorchis shared identical mass spectra profiles

We have developed a sensitive LC-MS/MS-based protocol to identify new steroids-derived molecules not only in extracts of helminth parasites [3,4], but also from experimental infected rodents [4] and naturally-infected humans [5]. Extracts obtained from *F. hepatica* adult worms were analyzed in order to provide insights related to their composition and complexity.

Comparing data obtained for *O. viverrini* with *O. felineus* we observed that both these liver flukes displayed highly similar mass spectra (MS) and shared most peaks detected (indicated in grey in Fig 1) which were attributed to oxysterol-like metabolites, e.g. mass/charge (m/z) 356, 307, bile acids in oxidized form, e.g. m/z 443, 479, 488 and DNA-adducts, e.g. m/z 599, 639, 667 [3,4].

**Fig 1.**
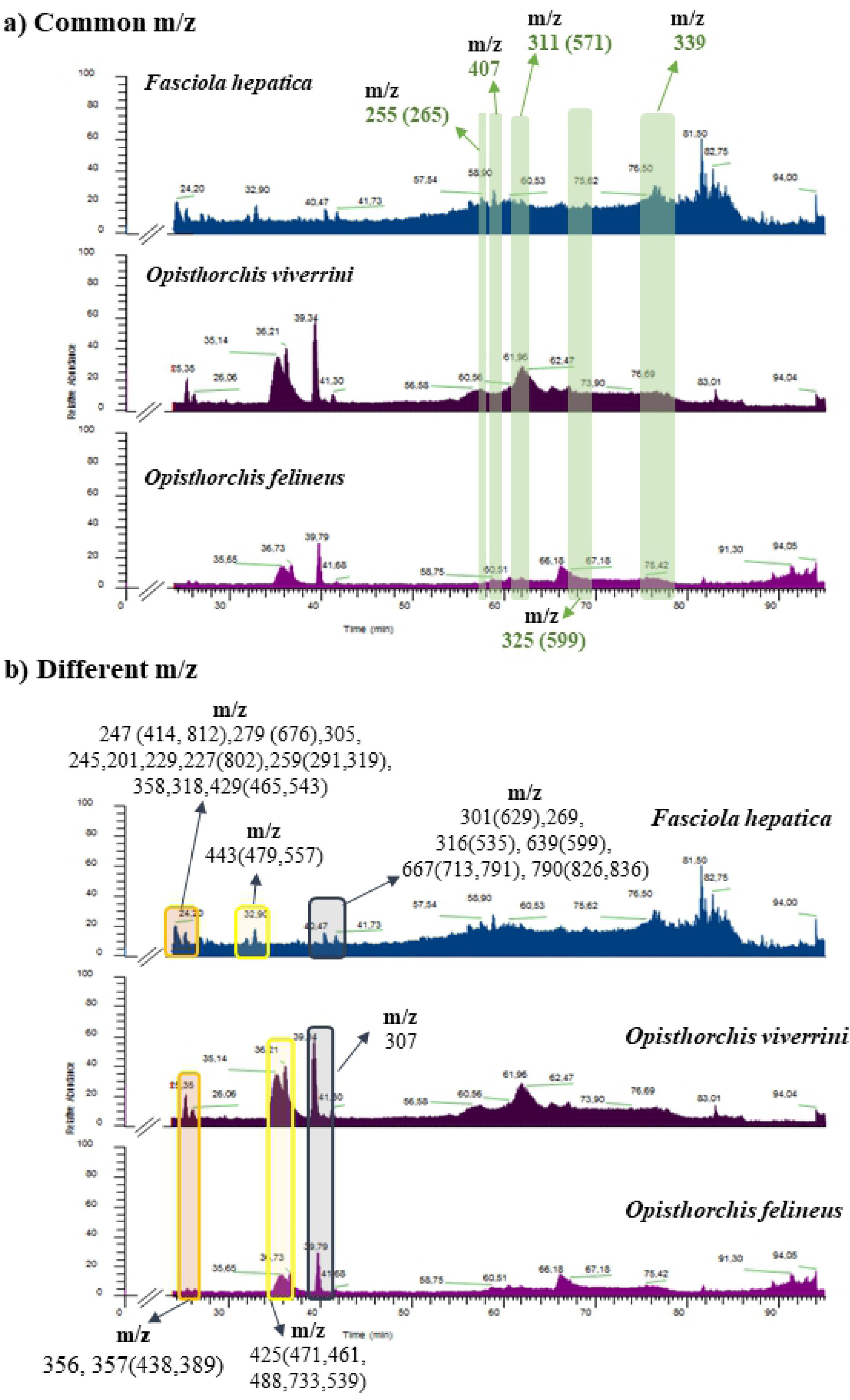
Comparison of mass spectral profiles obtained for *Fasciola hepatica* and *Opisthorchis* spp. Panel A, common m/z between the three liver flukes; panel B, major differences among the liver flukes.

### *F. hepatica* extracts exhibited striking differences to those of the *Opisthorchis* species

Notable differences were apparent among the MS profiles of *F. hepatica* and the *Opisthorchis* species. Most of compounds present in both *Opisthorchis* species were absent from *F. hepatica*, specifically m/z 356, 357, 425 and 307. Remarkably, these specific compounds were attributed to be oxysterols with ability to react with host DNA as described [3]. The MS profile of *F. hepatica* was much more complex than those obtained for *Opisthorchis* spp. (Fig 1). The major differences were observed at retention intervals of approximately 24, 32, and 40 min – as indicated in orange, yellow and blue, respectively, on the chromatographs (Fig 1). On these retention times, *F. hepatica* showed greater number of compounds in comparison to those observed on *Opisthorchis* species (Fig 1 and Table 2). Remarkably, most of these compounds were detected only in *F. hepatica* extracts (Table 2).

**Table 2.**
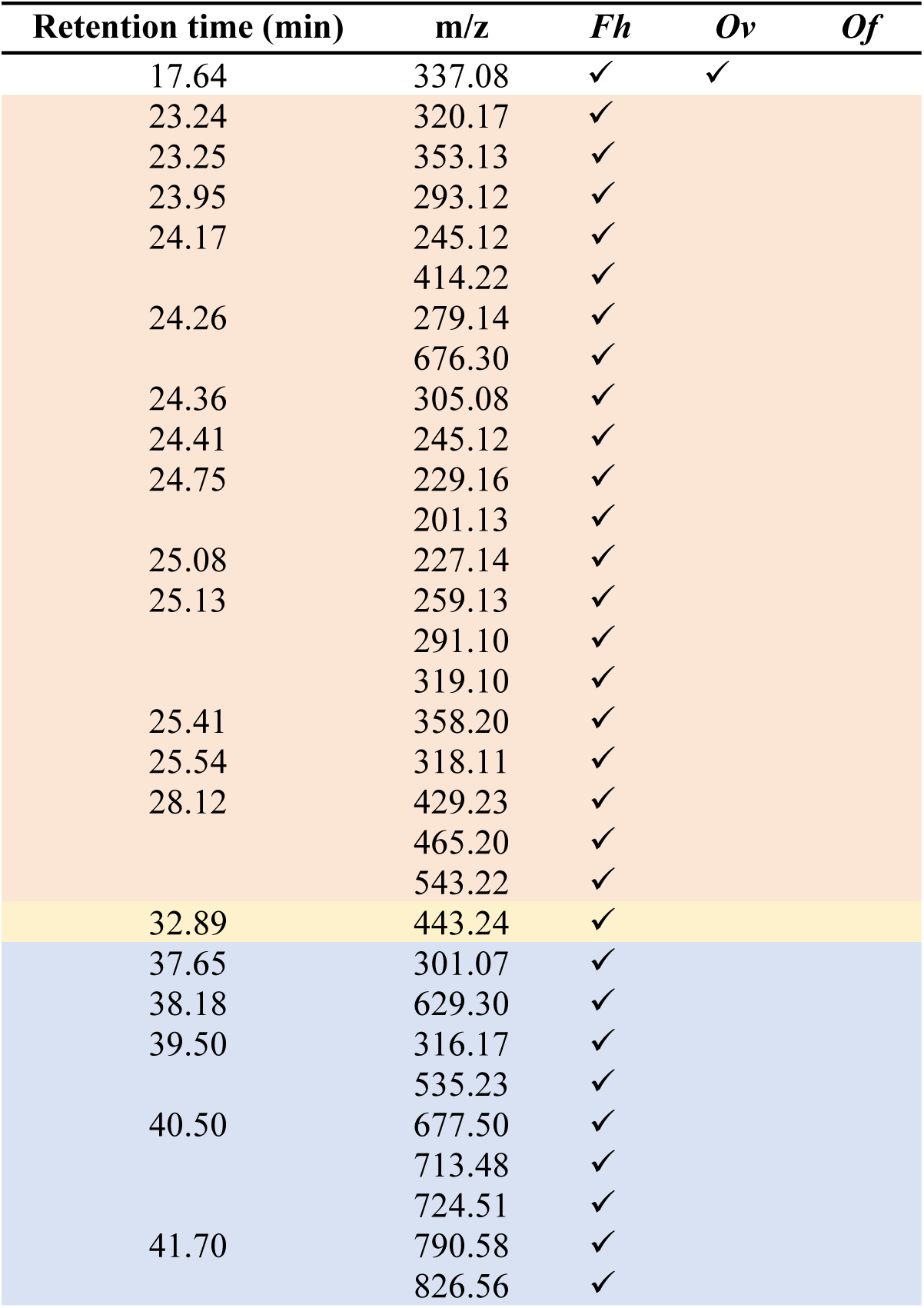
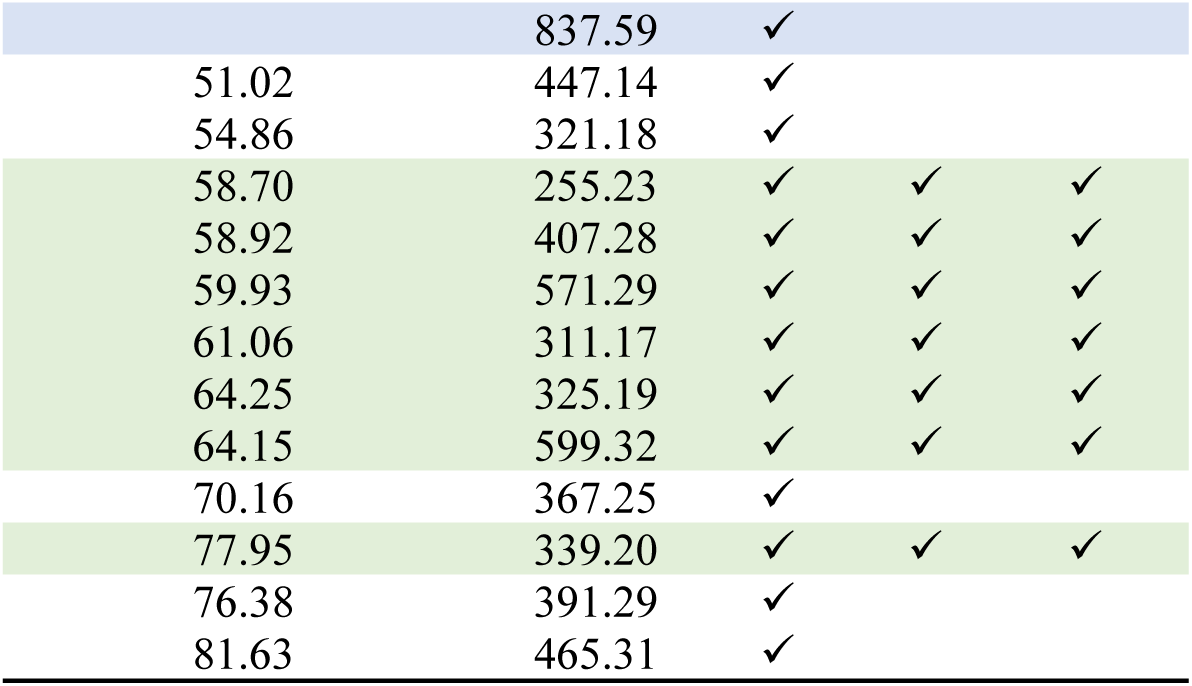
Comparison of mass/charge (m/z) obtained for *Fasciola hepatica* during this study with *Opisthorchis* spp. previously reported for *O. viverrini* [3,4] and *O. felineus* [3]. The structures of common m/z (signed at green) are depicted on S1 Table.

These findings suggested that metabolic processes of *F. hepatica were* notably distinct to those of *Opisthorchis* spp. Unlike *Opisthorchis, F. hepatica* displayed more compounds with elevated m/z (between 600 and 800), mostly between retention interval of 38 to 42 min (Table 2), which might suggest that they are more complex than the majority of those detected on *Opisthorchis* spp. This might be explained by the fact that *F. hepatica* juvenile parasites follow a different route to reach the biliary tree that involves ingestion and digestion of host tissues while traversing the intestinal wall, migrating through the peritoneal cavity and penetrating the liver Glisson’s capsule (S1A Fig). Moreover, during their liver migration the parasites ingest parenchymal and other cells [20] and these compounds might be a result of the complex metabolic pathways and profiles of these ingested tissues.

Nonetheless, *F. hepatica* and *Opisthorchis* spp. shared several common compounds at retention interval of 58-64 min (signed by green in Fig 1 and Table 2). These compounds have been ascribed previously to oxysterol-like metabolite (e.g. m/z 325), bile acids (e.g. m/z 571) and as well as DNA-adducts (m/z 599) [4]. This was expected since these parasites all reside within the bile ducts and fascioliasis can induce DNA damage [10,17]. To reiterate, however, these were fewer of these compounds in *F. hepatica* compared to *Opisthorchis* spp. Indeed, we posit that fewer oxysterol-like metabolites might (partially) explain why definitive carcinogenic potential has not been ascribed to ruminant or human fascioliasis (S1B Fig).

## Discussion

Chronic infection with *Fasciola* spp. and *Opisthorchis* spp. results in comparable pathology in their definitive hosts including fibrosis, hyperplasia and biliary stasis [3,10,21–23]. However, an association between fascioliasis and cancer remains controversial and not definitely established [10]. Thus, we decided to investigate extracts of adult worms of *F. hepatica* and compare with data previously obtained for *Opisthorchis* spp. We aimed to address the following questions: 1) does *F. hepatica* synthesize and excrete metabolites that promote direct damage on host DNA, and 2) if *F. hepatica* induces DNA damage, why has fascioliasis not been associated with liver cancer in ungulates or indeed humans? The MS profile of *F*. hepatica was found to be far more complex, showing an elevated number of compounds with an elevated m/z rather than *Opisthorchis* spp. This suggested that metabolic process that occur in *F. hepatica* are dissimilar to those in *Opisthorchis* spp.

The LC-MS/MS analysis also revealed a great diversity of compounds with different m/z. This diversity might reflect fragmentation of a number of compounds, detected here as lower m/z fragments of other compounds. However, we cannot conclude that these are not novel compounds. In addition, some of these compounds might be precursors of known compounds recorded previously [3,4]. Compounds of *F. hepatica* might be related to the different migratory route of the parasite to the biliary tree. Unlike *Opisthorchis* spp., newly excysted juveniles of *F. hepatica* exit the lumen of the small intestine, transverse the intestinal wall and migrate through the abdominal cavity to the Glisson’s capsule of the liver [20,24]. This parasite might deploy more complex biochemical processes and secretions, including the secretion of cathepsins [25–27] to accomplish this elaborate organ and tissue migration. The juvenile *F. hepatica* infects the liver by directly penetrating the Glisson’s capsule from the abdominal cavity, and thereafter burrows through the hepatic parenchyma to the bile ducts where it eventually matures into the egg-laying adult worm [20]. Components detected in the extracts of *F. hepatica* might be related with digestion of host tissues including blood such as hemoglobin, albumin and immunoglobin to support reproductive process including synthesis of eggs [20]. This might not only explain the complex MS profile but also the compounds with elevated m/z as well as lower m/z that could be associated with free amino acids. On other hand, most of the compounds observed from 23 to 57 minutes were specific of *F. hepatica*, i.e. not present in *Opisthorchis*. Juvenile *Opisthorchis* flukes ascend from the duodenum directly into the lumen of biliary tree [23,28].

Glycocholic acid in the mammalian small intestine triggers the excystment of the metacercaria and emergence of *F. hepatica* juvenile flukes stimulating the exit of the parasite from the gut lumen and its migration to the abdominal cavity. Intriguingly, the juvenile *F. hepatica* did not survive in bile-containing solutions whereas the adult fluke resides in the bile ducts, bathed in bile [29]. Differences in the nature of the juvenile versus adult tegument of *F. hepatica* and the selectivity and the permeability of glycocalyx of the tegument may underpin these stage specific differences [29]. The complexity of the tegument, a complex metabolically active and highly glycosylated biological matrix [30] might also underpin complexity of *F. hepatica* MS profile and its components.

Both *F. hepatica* and the two *Opisthorchis* species shared some identical compounds that were previously attributed to oxysterol-like metabolites, bile acids and DNA-adducts. This is feasible since all three flukes live within the biliary tree. There is evidence that *F. hepatica* induces DNA damage through the action of mutational-mediators [9,31]. The presence of DNA adducts in tissue does not necessarily imply a specific tumorigenic risk for the host tissue. Other factors such as DNA repair and cell proliferation key roles players in determining the overall carcinogenic risk [32]. An association between fascioliasis and cancer has only been suggested from *in vitro* studies and, thus far, there have not been satisfactory reports of human cases of bile duct cancer due to chronic infection with *F. hepatica* [10,14,33–35]. Therefore, there is a lack of cogent evidence that relate fascioliasis with cancer [10]. By contrast, a number of reports posit opposing effects, i.e. tumor growth stimulation and inhibition. Tumor growth stimulation and proliferation of hepatocytes has been observed during acute phase of infection where larval flukes migrate through the parenchyma of the liver and provoke marked inflammation [12,17]. In turn, the chronic inflammation increases oxidative stress that can overwhelm antioxidant system homeostasis to dampen reactive oxygen species and consequent oxidative modification of lipids, nucleic acids and proteins [9]. Like fascioliasis, opisthorchiasis is characterized by elevated oxidative stress and altered the antioxidant systems [9,11]. Tumor inhibition has been noted during the chronic phase of fascioliasis that may dampen the liver metabolizing activity [12]. We also documented that infection with *O. felineus* induces BilIN. The consonance of findings that the presence of new metabolites and of BilIN-1 and BilIN-2 indicates that *O. felineus* infection induces neoplastic transformation of cholangiocytes and can be expected to promote growth of biliary cancers [3]. Whereas acute *F. hepatica* infection may increase the metabolizing enzymes in liver and thus increase the activation of exogenous carcinogens [22], chronic infection may reduce hepatic metabolizing activity [12]. It is noteworthy that chronic infection with *F. hepatica* in a rat model suppressed *N*-nitrosodimethyldiamine-induced carcinogenesis, suggesting a parasite-induced inhibition of carcinogenesis in the liver of rodents experimentally infected with *F. hepatica* [17]. All these hypotheses require further investigation and experimental validation.

*Fasciola hepatica* displayed more complex mass spectra profile that the *Opisthorchis* species and several specific compounds that might be related to its complex route of migration to the biliary tract. Nonetheless, *F. hepatica* shared several compounds with *Opisthorchis*, which are related to oxysterols, bile acids and DNA-adducts. The presence of only a few common compounds might explain why fascioliasis has not been causally linked with liver cancer. On the other hand, it has been shown that *F. hepatica* could suppress reactivity of liver carcinogens.

## Acknowledgments

This work was financed by FEDER - Fundo Europeu de Desenvolvimento Regional funds through the COMPETE 2020 - Operacional Programme for Competitiveness and Internationalisation (POCI), Portugal 2020, and by Portuguese funds through FCT - Fundação para a Ciência e a Tecnologia, in the framework of the project, Institute for Research and Innovation in Health Sciences” (POCI-01-0145-FEDER-007274). The FCT and FEDER (European Union) also supported these studies through project number IF/00092/2014/CP1255/CT0004. NV thanks FCT by IF position, Fundação Manuel António da Mota (FMAM, Portugal) and Pfizer Portugal by support Nuno Vale Lab. JMCC thanks FCT for Pest-OE/AGR/UI0211/2011 and Strategic Project UI211. PJB gratefully acknowledges support from award CA164719, National Cancer Institute, National Institutes of Health (NIH). MYP and VAM acknowledge the support from the Russian Science Foundation, project number 18-15-00098. The contents of this report are solely the responsibility of the authors and do not necessarily represent the official views of the FCT, FMAM, Pfizer Portugal or the NIH.

## Declarations of conflicting interests

The authors declare no conflicts of interest with respect to the research, authorship, and/or publication of this article.

## Supporting information

**S1 Table.** Structures of m/z common to *Fasciola hepatica* and *Opisthorchis* species.

**S1A Fig. Different routes that liver flukes undergo to reach the biliary tree**. *F. hepatica* (signed at blue) transverses the intestinal wall and migrates through peritoneum to the Glisson’s capsule of the liver, perforate the capsule enters the liver parenchyma and migrates to the biliary tree. In contrast, Opisthorchis spp. juveniles pass through the stomach to the duodenum with ingested fish, after which they ascend into the biliary tract through the ampulla de Vater (signed at yellow). This might constitute the major reason for complexity of mass spectra profile of *F. hepatica*.

**S1B Fig. Adult liver flukes *O. viverrini* and *O. felineus* produces oxysterol-like metabolites that interact with host chromosomal DNA to form DNA-adducts and forms of biliary intraepithelial neoplasia that conducive to cholangiocarcinoma.** *F. hepatica* also elaborates oxysterol-like metabolites, but at much lower number, which might be explain, at least in part, why infection with this parasite fails to induce malignancy.

## Graphical Abstract

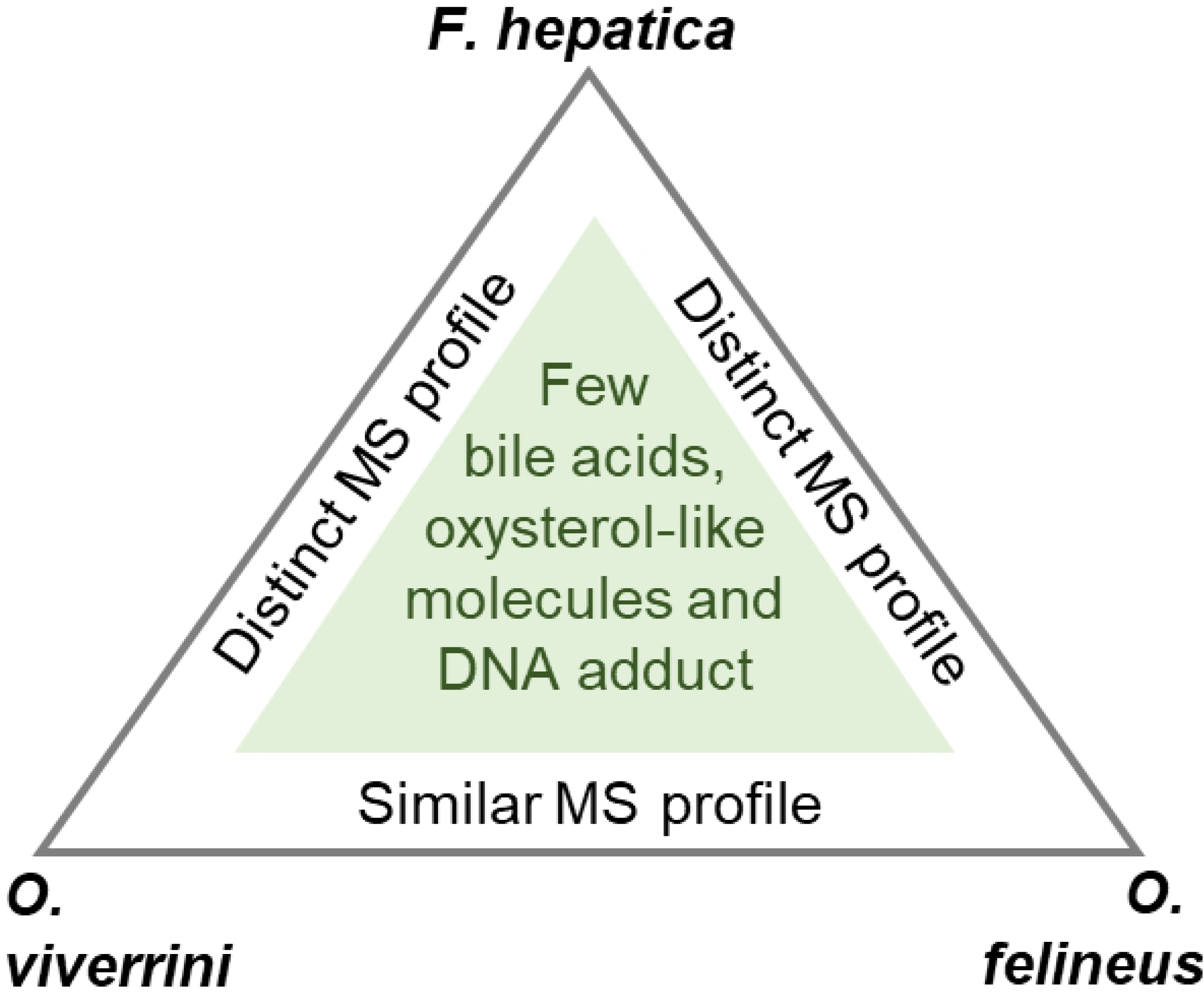

